# AGouTI - flexible Annotation of Genomic and Transcriptomic Intervals

**DOI:** 10.1101/2022.11.13.516331

**Authors:** Jan G. Kosiński, Marek Żywicki

**Affiliations:** Department of Computational Biology, Institute of Molecular Biology and Biotechnology, Faculty of Biology, Adam Mickiewicz University in Poznań, 61-614 Poznań, Poland

## Abstract

**Summary:** The recent development of high-throughput workflows in genomics and transcriptomics revealed that efficient annotation is essential for effective data processing and analysis. Although a variety of tools dedicated to this purpose is available, their functionality is limited. Here, we present *AGouTI* – a universal tool for flexible annotation of any genomic or transcriptomic coordinates using known genomic features deposited in different publicly available databases in the form of GTF or GFF files. In contrast to currently available tools, *AGouTI* is designed to provide a flexible selection of genomic features overlapping or adjacent to annotated intervals, can be used on custom column-based text files obtained from different data analysis pipelines, and supports operations on transcriptomic coordinate systems.

*Availability and Implementation:* AGouTI was implemented using Python 3 and is freely available on GitHub (https://github.com/zywicki-lab/agouti), from the Python Package Index (https://pypi.org/project/AGouTI/) or Anaconda Cloud (https://anaconda.org/bioconda/agouti). We also provide a Galaxy wrapper available from the Galaxy Tool Shed (https://toolshed.g2.bx.psu.edu/view/janktoolshed/agouti/c204da8f836d).

*Supplementary information:* Supplementary Data are available at Publisher site online. Files for replicating the use-case scenario described in Supplementary Data are available at Zenodo: https://doi.org/10.5281/zenodo.7317210.

*Contact:*

## Introduction

In recent years, high-throughput techniques have become a standard research tool, providing a massive increase in the amount of data produced daily in research laboratories across the globe and stored in public databases. Most often, high-throughput approaches are used for large-scale characterization of whole genomes, transcriptomes, or proteomes. As a result, the localization of specific biological sequences, structures, and other distinctive features is obtained and stored using a coordinate-based system in one of the standard formats (BED, GFF/GTF, VCF, and others) or custom tables.

It is of particular interest for scientists to obtain comprehensive information about the regions of interest in the context of known genomic features such as genes, regulatory regions, UTRs, or CDSs to infer their biological role or function. Numerous tools have been developed to help scientists achieve this goal. Two most widely used include the *BEDTools* suite (Quinlan, Hall 2010) - a powerful toolset for genome arithmetic - with such commands as *intersect, annotate*, and *closest*; and the *BEDOPS* suite (Neph et al. 2012) with *closest-features* and *bedmap* programs. Many other pipelines and tools designed for that, or similar purpose exist, e.g., *annotatePeaks*.*pl* from *Homer* (Heinz et al. 2010) or *AnnotateGenomicRegions* (Zammataro et al. 2014).

Although all those tools are widely used for annotating genomic coordinates, none can work with transcriptomic data represented as transcript-based coordinates or custom text tables. To fill this gap, we have developed *AGouTI* - a tool for flexible Annotation of Genomic and Transcriptomic Intervals, designed to annotate any genomic or transcriptomic coordinates using user-selected features from genome annotations stored in GTF or GFF files.

## Design and Implementation

The workflow of *AGouTI* is divided into two major steps (Figure 1). First, the SQLite database is built based on genome annotations provided in a GTF or GFF file (*agouti create_db* command). The database enables fast and memory-efficient access to genomic features during annotation. Since it is written on a hard drive, it can be used in multiple projects, significantly reducing the analysis time.

**Figure 1.**
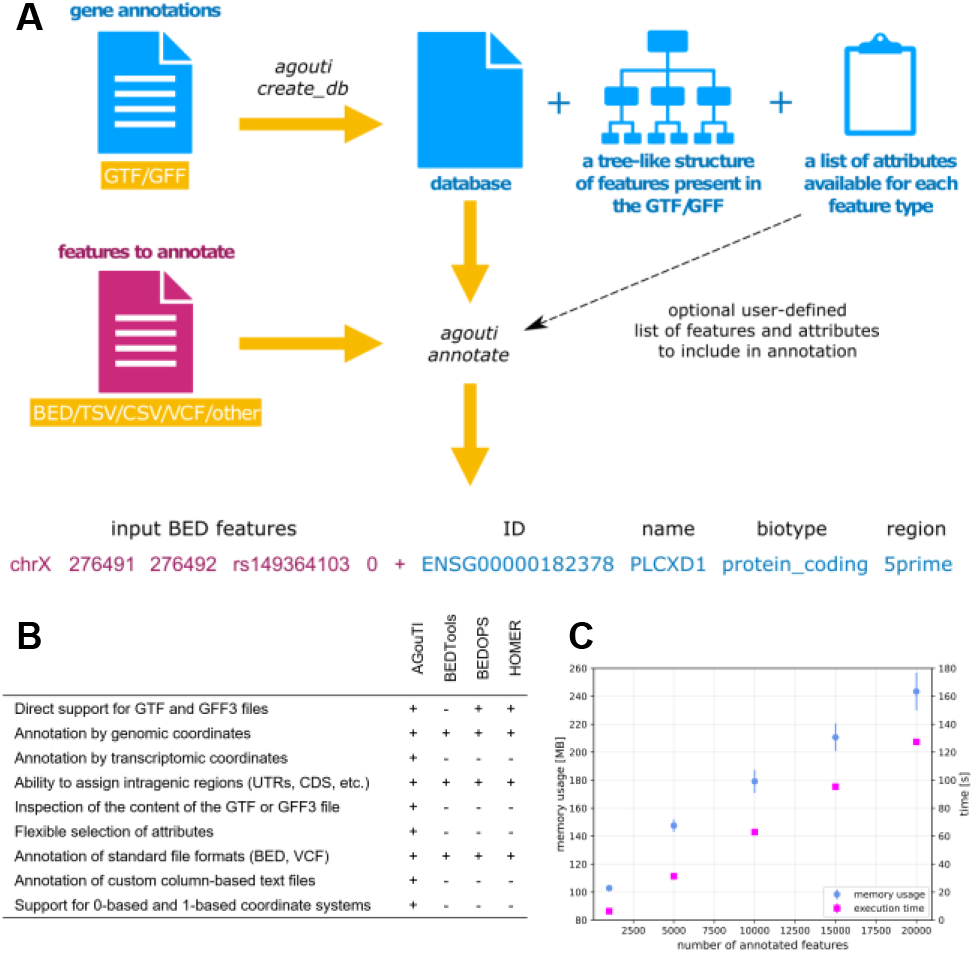
Overview of *AGouTI*. (A) Schematic representation of the annotation process; (B) comparison of *AGouTI* features with other similar tools; (C) time and memory complexity of *agouti annotate* process depending on the size of the annotated dataset. Error bars represent the SD for 10 repetitions.

Once the database is created, the *agouti annotate* command can be used to initiate the annotation process. This step utilizes the previously created database to identify the features and attributes overlapping with genomic or transcriptomic intervals from the submitted column-based text file (usually BED or VCF). The annotation is appended to the end of each line as additional columns. The results are displayed on the standard output in a tabular and self-explanatory format.

*AGouTI* was implemented using Python 3. To provide comprehensive compatibility with the GTF/GFF format, the processing of genome annotation files has been implemented using the *gffutils* python module (Dale 2014).

## Features

*AGouTI* provides several features not available in other annotation software (Figure 1B). The significant advantage is the ability to annotate regions defined by genome-based and transcript-based coordinates (defined as transcript ID and positions within the transcript). In the latter case, the annotation includes the identification of the parental gene (using transcript ID) and all the attributes assigned to child records in the GFF/GTF file, including encoded protein and overlapping intragenic features, such as CDS, UTRs, and others. In the case of annotation of genome-based coordinates, overlapping intragenic features can also be assigned, as well as the closest adjacent genes. Additionally, *AGouTI* calculates the relative position within the overlapping or adjacent feature, reporting it as *upstream, downstream, 3’* or *5’ part, middle, whole*, or *full* (see Supplementary Data and the AGouTI documentation on GitHub for details).

The second advantage of *AGouTI* is the ability to annotate any column-based text files containing at least information about feature ID, reference (chromosome or transcript ID), start and end positions located in specific columns. The columns containing that information and the column separator can be specified using appropriate program options. The overlapping genomic features are appended to each line as additional columns without alteration of the original data format.

*AGouTI* has been designed to provide the highest possible flexibility in the annotation process. It is fulfilled by extensive control over selecting features and their attributes to be included in the annotation. First, the GFF/GTF file is inspected during the building of the database, and a comprehensive list of all feature types and assigned attributes is presented to the user (Supplementary Figure 2). Next, while performing the annotation, the user can assign all available information (default) or select only those relevant to the annotated data (see a use-case scenario in Supplementary Data). It is also possible to decide whether the annotation should be performed on the gene or transcript level. In case of multiple genes or transcripts overlapping a given interval, all hits are returned as individual lines.

Other essential features of AGouTI include the ability to use both 0 or 1-based coordinate systems and to select features entirely or partially overlapping annotated intervals. *AGouTI* can also provide the statistics of annotated features and attributes, e.g., counts of gene or transcript biotypes and CDS/UTR distribution, depending on a set of information selected for annotation (Supplementary Figure 3). Although *AGouTI* contains many options allowing for extensive customization of the annotation process, it is straightforward to use. In the case of using the default options (input file containing genomic coordinates in BED format, selection of all available features and attributes), it requires specifying only the input files. An example usecase scenario is described in the Supplementary Data. The installation process is also user-friendly – the packages are available via Python Package Index and Anaconda Cloud. Despite the implementation of multiple computationally demanding features, AGouTI remains very efficient. Both annotation (Figure 1C) and database building (Supplementary Figure 1) are characterized by linear time and memory complexity. For more details on benchmarking results, see Supplementary Data.

## Conclusion

*AGouTI* is an efficient and highly customizable tool for annotating genomic and transcriptomic intervals. The unique features implemented in the software make it the first tool to efficiently annotate results from the vast majority of data analysis tools and pipelines in genomics and transcriptomics.

## Data availability

There are no new data associated with this article.

