## Supplementary Data File for "AGouTI - flexible Annotation of Genomic and Transcriptomic Intervals"

### Supplementary Information

**Jan G. Kosiński, Marek Żywicki**

**November 13, 2022**

### Benchmarking the *AGouTI pipeline*

In order to compare the processing speed of AGouTI in genomic mode with its most popular counterpart - *bedtools intersect*- we have annotated 104 573 transcriptions start sites (TSS) from Brassica napus. Coordinates of TSS were downloaded from the Ensembl Plants database (Brassica napus genes AST_PRJEB5043_v1; *http://plants.ensembl.org/index.html*; release 53) (Cunningham et al. 2022) using the BioMart tool and converted to BED format with AWK. The *Brassica napus* genome annotation was downloaded in GTF format from the Ensembl Plants database (release 53). The annotation using both tested tools was performed using default settings.
The tests revealed that *AGouTI* is slower than *bedtools intersect*, reaching a processing speed of 288 against 10 000 annotated TSS per second, respectively. The result was, however, expected, since the feature selection, multi-level feature annotation, and estimation of intragenic regions implemented in *AGouTI* are computationally complex routines providing unique functionality by the cost of computing time. Moreover, *AGouTI* provides a combined functionality of two different tools from the *bedtools* suite: *intersect* and *closest*.
To get more insight into the time and memory complexity of the AGouTI algorithm in the transcriptomic mode, we decided to use the most computationally demanding scenario employing the unique features of AGouTI. Namely, in the command invocation, we have used options dedicated to handling custom file format, calculating the intragenic location, and generating summary statistics. A detailed procedure, including specific commands, has been described in the below use-case scenario. The miRNA binding sites obtained using psRNATarget (see the description below) were multiplied to obtain up to 20 000 records to measure *AGouTI* running times and memory footprint on datasets having diverse sizes. In order to simulate the real-life use of *AGouTI*, benchmark runs were performed on a laptop computer (MacBook Air M1 laptop (2020) with 16GB RAM) and calculated as an average of 10 independent command invocations. Measurements were made using the GNU implementation of the time utility (https://www.gnu.org/software/time/).
The benchmark revealed that in both steps – database build and annotation, the time and memory usage have linear complexity with a relatively small footprint, allowing efficient use of the tool on standard computer setups. Compared to genomic mode, the processing speed of the *agouti annotate* with the above-described computationally-demanding combination of features dropped from 288 to 160 intervals per second. The detailed benchmarking results for the annotation process have been presented in the main manuscript (Figure 1C) and for database creation in Supplementary Figure 1.

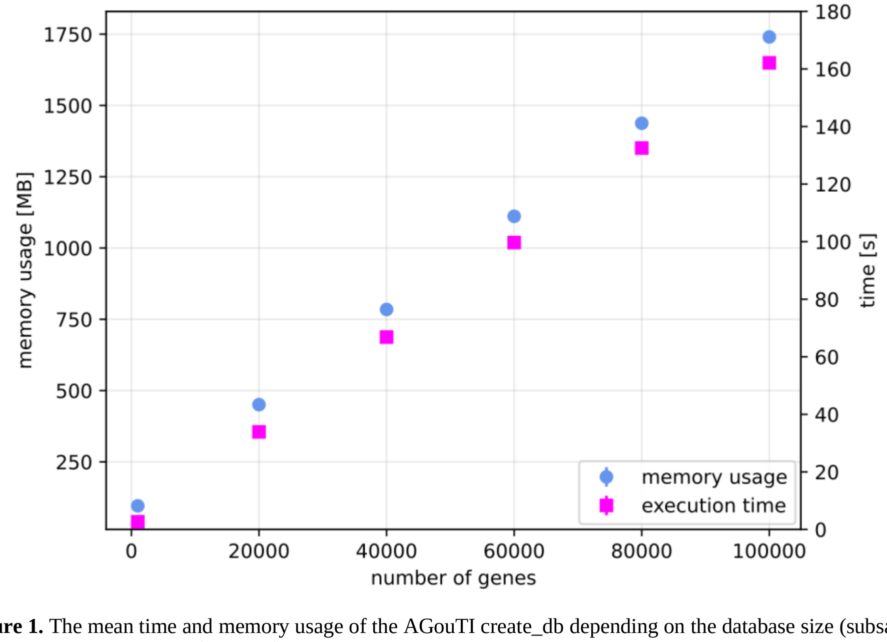

**Supplementary Figure 1.** The mean time and memory usage of the AGouTI create_db depending on the database size (subsampled 1,20,40,60,80 and 100 thousand genes along with their subfeatures from the *Brassica napus* genes AST_PRJEB5043_v1). Each run has been repeated independently 10 times.

### Example use-case scenario

As the use-case of the AGouTI pipeline, we describe the annotation of miRNA target regions predicted in mRNA sequences of *Brassica napus* with the psRNATarget tool (https://www.zhaolab.org/psRNATarget/home) (Xinbin Dai et al. 2018, 2011, 2011). The output of the psRNATarget is a tab-separated text file containing information about predicted interaction sites, including their positions within the mRNA sequence. The input files necessary for completing the below-described workflow are available on Zenodo (DOI: 10.5281/zenodo.7317210; https://doi.org/10.5281/zenodo.7317210).

For prediction of the miRNA binding sites, the online version of *psRNATarget* has been used with default settings (as of 07/07/2022) on the input consisting of 92 *Brassica napus* miRNA sequences from the miRBase database (release 21, June 2014). *Brassica napus* transcriptome has been downloaded from the Ensembl Plants database (release 53) and restricted to the first 40 000 transcript sequences to limit the volume of the example dataset. The results have been downloaded and saved as the *psRNATarget_bnapus.tsv*. The structure of the file has been presented in Supplementary Table 1.

For annotation of the predicted miRNA binding sites in mRNA, we have downloaded the *Brassica napus* genome annotation in GTF format from the Ensembl Plants database (release 53; *Brassica_napus.AST_PRJEB5043_v1.53.gtf.gz*).

The first step of the AGouTI pipeline is to create a dedicated database based on the annotation file. The user can easily do this with the following command:

```
agouti create_db -f GTF -a Brassica_napus.AST_PRJEB5043_v1.53.gtf.gz -d bnapus.db
```

**Average execution time:** ~169s
**Maximum memory usage:** ~1796MB
(tested on MacBook Air M1 2020, with 16GB RAM, calculated as an average of 10 independent command invocations)

As a result, several files have been created, including the SQLite database (*bnapus.db*) and the file describing the structure, features, and attributes of the GTF file (*bnapus.db.database.structure.txt*; Supplementary Figure 2).

```
Attributes available for each feature type:

gene: gene_id, gene_biotype, gene_source, gene_name
transcript: gene_biotype, gene_source, gene_name, transcript_biotype, transcript_id, gene_id,
transcript_name, transcript_source
exon: gene_source, gene_biotype, gene_name, transcript_biotype, exon_id, transcript_id, gene_id,
exon_number, transcript_name, transcript_source
cds: gene_source, gene_biotype, gene_name, transcript_biotype, transcript_id, protein_id, gene_id,
exon_number, transcript_name, transcript_source
start_codon: gene_source, gene_biotype, gene_name, transcript_biotype, transcript_id, gene_id,
exon_number, transcript_name, transcript_source
stop_codon: gene_source, gene_biotype, gene_name, transcript_biotype, transcript_id, gene_id,
exon_number, transcript_name, transcript_source
five_prime_utr: gene_biotype, gene_source, gene_name, transcript_biotype, transcript_id, gene_id,
transcript_name, transcript_source
three_prime_utr: gene_biotype, gene_source, gene_name, transcript_biotype, transcript_id, gene_id,
transcript_name, transcript_source

gene
└── transcript
    ├── stop_codon
    ├── three_prime_utr
    ├── start_codon
    ├── five_prime_utr
    ├── exon
    └── cds
```

**Supplementary Figure 2.** The content of the *bnapus.db.database.structure.txt* file describing the input annotation file's structure and the attributes of each feature type available for selection during annotation.

Once the database is prepared, it is possible to run the annotation procedure. AGouTI requires each interval to have a unique ID. The *psRNATarget_bnapus.tsv* file doesn't have a column with such an ID, meaning it needs to be added before proceeding with the AGouTI pipeline. This can be done simply by adding in any spreadsheet software (e.g., Microsoft Excel) a column with unique ascending numbers  or by running the following awk command:

```
awk 'NR>2{print(NR"-"$0)} NR <=2 {print}' psRNATarget_bnapus.tsv > psRNATarget_bnapus_with_ID.tsv
```

**Tip!** The value of 2 in the expressions "NR>2" and "NR <= 2" of the command corresponds to the number of header lines present in the file, which should be excluded from the process.

The annotation process can be run using:

```
agouti annotate -d bnapus.db -i psRNATarget_bnapus_with_ID.tsv -n 2 --custom 1,2,7,8 -r -t --statistics -p
"\t" -a gene_biotype,transcript_biotype > agouti_miRNA-targets.tsv 2> agouti_statistics.txt
```

**Average execution time:** ~73s
**Maximum memory usage:** ~207MB
(12137 annotated features (lines), tested on MacBook Air M1 2020, with 16GB RAM, calculated as an average of 10 independent command invocations)

The annotated (input) *psRNATarget* result file (*psRNATarget_bnapus_with_ID.tsv*) is a TSV file with the custom column order (see Supplementary Table 1). Therefore, the user needs to specify (the --*custom* option) the column numbers containing feature id (1$^{st}$ column), transcript (reference sequence) id (2$^{nd}$), start (7$^{th}$), and end (8$^{th}$) coordinates of the predicted interaction site. Column separator can be defined with the *-p* flag and the number of header lines with *-n*. In our file, coordinates are described using a transcript-based coordinate system, thus, the option *-t* is also necessary. Although the intragenic regions defined in the GTF file, overlapping with annotated regions, are returned by default, the user can also decide to calculate the intragenic location of the entry from the input file by adding the *-r* flag. Possible values returned by this option are:

- **5 prime** - when the annotated interval starts within the first quarter of gene or transcript and ends in the first half
- **middle** - interval starts and ends within the second and third quarter, respectively
- **3 prime** - interval starts within the third quarter and ends in the last one
- **whole** - interval starts within the first quarter and ends within the last one. The annotated interval size **does not exceed** 90% of the transcript or gene length.
- **full** - interval starts within the first quarter and ends within the last one. The annotated interval size **exceeds** 90% of the transcript or gene length.
- **upstream** – interval is located upstream to the gene or transcript
- **downstream** – interval is located downstream to the gene or transcript.

Furthermore, the user can decide that the only valuable attributes to include in the annotation are '*gene_biotype*' and '*transcript_biotype*' by applying the *-a* flag. The --*statistics* option can be used to generate an additional summary displayed on the stderr by default (see *agouti_statistics.txt* and Supplementary Figure 3). The main output of the *agouti annotation* command is displayed on the stdout (see *agouti_miRNA-targets.tsv* and Supplementary Table 2).

```
###################
##STATISTICS
###################
# statistics for the 'cds' column: 'y' - 9963; '.' - 2173
# statistics for the 'five_prime_utr' column: '.' - 11285; 'y' - 851
# statistics for the 'three_prime_utr' column: '.' - 10703; 'y' - 1433
# statistics for the 'gene_biotype' column: 'protein_coding' - 12136
# statistics for the 'transcript_biotype' column: 'protein_coding' - 12136
# statistics for the 'feature_region' column: 'middle' - 5734; '3 prime' - 3449;
'5 prime' - 2948; 'other' - 5
```

**Supplementary Figure 3.** Statistics generated by the AGouTI pipeline based on the output file.

**Supplementary Table 1.** The first rows of the *psRNATarget_bnapus.tsv* file as a schematic representation of the custom input file format.

| #Please import the downloaded file into Microsoft Excel or other spreadsheet software | | | | | | | | | | | | | |
|---|---|---|---|---|---|---|---|---|---|---|---|---|---|
| **miRNA_Acc.** | Target_Acc. | Expectation | UPE$ | miRNA_start | miRNA_end | Target_start | Target_end | miRNA_aligned_fragment | alignment | Target_aligned_fragment | Inhibition | Target_Desc. | Multiplicity |
| **bna-miR156a** | CDY69014 | 0.0 | -1.0 | 1 | 21 | 682 | 702 | UGACAGAAGAGAGUGAGCACA | ::::::::::::::::::::: | UGUGCUCACUCUCUUCUGUCA | Cleavage | | 1 |
| **bna-miR156d** | CDY69014 | 0.0 | -1.0 | 1 | 20 | 683 | 702 | UGACAGAAGAGAGUGAGCAC | :::::::::::::::::::: | GUGCUCACUCUCUUCUGUCA | Cleavage | | 1 |
| **bna-miR156e** | CDY69014 | 0.0 | -1.0 | 1 | 20 | 683 | 702 | UGACAGAAGAGAGUGAGCAC | :::::::::::::::::::: | GUGCUCACUCUCUUCUGUCA | Cleavage | | 1 |
| **bna-miR156f** | CDY69014 | 0.0 | -1.0 | 1 | 20 | 683 | 702 | UGACAGAAGAGAGUGAGCAC | :::::::::::::::::::: | GUGCUCACUCUCUUCUGUCA | Cleavage | | 1 |
| **bna-miR171f** | CDY56248 | 0.0 | -1.0 | 1 | 21 | 774 | 794 | UGAUUGAGCCGCGCCAAUAUC | ::::::::::::::::::::: | GAUAUUGGCGCGGCUCAAUCA | Cleavage | | 1 |
| **bna-miR171f** | CDX95633 | 0.0 | -1.0 | 1 | 21 | 811 | 831 | UGAUUGAGCCGCGCCAAUAUC | ::::::::::::::::::::: | GAUAUUGGCGCGGCUCAAUCA | Cleavage | | 1 |
| **bna-miR171f** | CDY67934 | 0.0 | -1.0 | 1 | 21 | 799 | 819 | UGAUUGAGCCGCGCCAAUAUC | ::::::::::::::::::::: | GAUAUUGGCGCGGCUCAAUCA | Cleavage | | 1 |
| **bna-miR171f** | CDY63062 | 0.0 | -1.0 | 1 | 21 | 832 | 852 | UGAUUGAGCCGCGCCAAUAUC | ::::::::::::::::::::: | GAUAUUGGCGCGGCUCAAUCA | Cleavage | | 1 |

**Supplementary Table 2.** The first rows of the *psRNATarget_bnapus.tsv* file annotated with the *AGouTI* pipeline.

| #Please import the downloaded file into Microsoft Excel or other speadsheet software | | | | | | | | | | | | | | | | | | | | | | | | |
|---|---|---|---|---|---|---|---|---|---|---|---|---|---|---|---|---|---|---|---|---|---|---|---|---|
| **feature_id** | transcript_id | Expectation | UPE$ | miRNA_start | miRNA_end | feature_start | feature_end | miRNA_aligned_fragment | alignment | Target_aligned_fragment | Inhibition | Target_Desc. | Multiplicity | annotated_gene_id | annotated_featutype | annotated_chromosome | annotated_transcript_start | annotated_transcript_end | cds | five_prime_utr | three_prime_utr | gene_biotype | transcript_biotype | feature_region |
| **3-bna-miR156a** | CDY69014 | 0.0 | -1.0 | 1 | 21 | 682 | 702 | UGACAGAAGAGAGUGAGCACA | ::::::::::::::::::::: | UGUGCUCACUCUCUUCUGUCA | Cleavage | | 1 | GSBRNA2T00082020001 | transcript | LK038451 | 2498 | 3741 | y | . | . | protein_coding | protein_coding | middle |
| **4-bna-miR156d** | CDY69014 | 0.0 | -1.0 | 1 | 20 | 683 | 702 | UGACAGAAGAGAGUGAGCAC | :::::::::::::::::::: | GUGCUCACUCUCUUCUGUCA | Cleavage | | 1 | GSBRNA2T00082020001 | transcript | LK038451 | 2498 | 3741 | y | . | . | protein_coding | protein_coding | middle |
| **5-bna-miR156e** | CDY69014 | 0.0 | -1.0 | 1 | 20 | 683 | 702 | UGACAGAAGAGAGUGAGCAC | :::::::::::::::::::: | GUGCUCACUCUCUUCUGUCA | Cleavage | | 1 | GSBRNA2T00082020001 | transcript | LK038451 | 2498 | 3741 | y | . | . | protein_coding | protein_coding | middle |
| **6-bna-miR156f** | CDY69014 | 0.0 | -1.0 | 1 | 20 | 683 | 702 | UGACAGAAGAGAGUGAGCAC | :::::::::::::::::::: | GUGCUCACUCUCUUCUGUCA | Cleavage | | 1 | GSBRNA2T00082020001 | transcript | LK038451 | 2498 | 3741 | y | . | . | protein_coding | protein_coding | middle |
| **7-bna-miR171f** | CDY56248 | 0.0 | -1.0 | 1 | 21 | 774 | 794 | UGAUUGAGCCGCGCCAAUAUC | ::::::::::::::::::::: | GAUAUUGGCGCGGCUCAAUCA | Cleavage | | 1 | GSBRNA2T00018011001 | transcript | LK033471 | 43638 | 45443 | y | . | . | protein_coding | protein_coding | middle |
| **8-bna-miR171f** | CDX95633 | 0.0 | -1.0 | 1 | 21 | 811 | 831 | UGAUUGAGCCGCGCCAAUAUC | ::::::::::::::::::::: | GAUAUUGGCGCGGCUCAAUCA | Cleavage | | 1 | GSBRNA2T00101513001 | transcript | LK031890 | 361274 | 363227 | y | . | . | protein_coding | protein_coding | middle |
| **9-bna-miR171f** | CDY67934 | 0.0 | -1.0 | 1 | 21 | 799 | 819 | UGAUUGAGCCGCGCCAAUAUC | ::::::::::::::::::::: | GAUAUUGGCGCGGCUCAAUCA | Cleavage | | 1 | GSBRNA2T00067696001 | transcript | LK036669 | 3839 | 5672 | y | . | . | protein_coding | protein_coding | middle |
| **10-bna-miR171f** | CDY63062 | 0.0 | -1.0 | 1 | 21 | 832 | 852 | UGAUUGAGCCGCGCCAAUAUC | ::::::::::::::::::::: | GAUAUUGGCGCGGCUCAAUCA | Cleavage | | 1 | GSBRNA2T00037888001 | transcript | LK034239 | 20805 | 22668 | y | . | . | protein_coding | protein_coding | middle |
